# Low Intensity Vibration Restores Nuclear YAP Levels and Acute YAP Nuclear Shuttling in Mesenchymal Stem Cells Subjected to Simulated Microgravity

**DOI:** 10.1101/2020.05.04.076711

**Authors:** M Thompson, K Woods, J Newberg, JT Oxford, G Uzer

## Abstract

Reducing the bone deterioration that astronauts experience in microgravity requires countermeasures that can improve the effectiveness of rigorous and time-expensive exercise regimens under microgravity. The ability of low intensity vibrations (LIV) to activate force-responsive signaling pathways in cells suggests LIV as a potential countermeasure to improve cell responsiveness to subsequent mechanical challenge. Mechanoresponse of mesenchymal stem cells (MSC) which maintain bone-making osteoblasts is in part controlled by the “mechanotransducer” protein YAP (Yes-associated protein) which is shuttled into the nucleus in response cyto-mechanical forces. Here, using YAP nuclear shuttling as a measure of MSC mechanoresponse, we tested the effect of 72 hours of simulated microgravity (SMG) and daily LIV application (LIV_DT_) on the YAP nuclear entry driven by either acute LIV (LIV_AT_) or Lysophosphohaditic acid (LPA), applied at the end of the 72h period. We hypothesized that SMG-induced impairment of acute YAP nuclear entry will be alleviated by daily application of LIV_DT_. Results showed that while both acute LIV_AT_ and LPA treatments increased nuclear YAP entry by 50% and 87% over the basal levels in SMG-treated MSCs, nuclear YAP levels of all SMG groups were significantly lower than non-SMG controls. Daily dosing of LIV_DT_, applied in parallel to SMG, restored the SMG-driven decrease in basal nuclear YAP to control levels as well as increased the LPA-induced but not LIV_AT_-induced YAP nuclear entry over the non-LIV_DT_ treated, SMG only, counterparts. These cell level observations suggest that utilizing daily LIV treatments is a feasible countermeasure for increasing the YAP-mediated anabolic responsiveness of MSCs to subsequent mechanical challenge under SMG.

## Introduction

The musculoskeletal deterioration which astronauts experience on long-term space missions and the resulting increase of traumatic physical injury risk is in part due to the reduction of mechanical loading on the musculoskeleton ^1^. To alleviate the detrimental effects of unloading, astronauts undergo intensive regimens of running and resistance training in orbit ^2^. Despite these efforts, astronauts lose an average bone density of 1% for each month they spend in space ^3^. This loss necessitates new non-pharmacologic therapies in addition to exercise to keep bones healthy during long-term space missions. In bone, tissue level response to mechanical challenge is in part regulated by osteoblasts and osteocytes ^4^. Both osteoblasts and osteocytes in turn share a common progenitor: the mesenchymal stem cell (MSC). Therefore, the growth and differentiation of MCSs in response to mechanical stimulation is required for the maintenance and repair of bone ^5^. It is for this reason that the MSCs are a potential target for mechanical therapies aiming to alleviate bone loss in astronauts, injured service personnel with long periods of bedrest, and physically inactive aged individuals ^6^.

To maintain healthy bone making cell populations, MSCs rely on environmental mechanical signals inside the bone marrow niches and near bone surfaces. While the exact characteristics of the mechanical environment in which MSCs exist remains to be quantified, it is known that during habitual activities, our bones are subjected to combinations of complex loads including strain, fluid shear, and acceleration, each of which is inseparable ^7^. For example, during moderate running, cortical bone can experience strains up to 2000µε ^8,9^, which also generates coupled fluid flow within canaliculi of up to 100µm/s ^10^. The interior of bone is filled with bone marrow with viscosities in the range of 400-800cP ^11^. During moderate running, tibial accelerations are within the 2-5g range ^12^ (1g = 9.81 m/s^2^), creating a complex loading at the bone-marrow interface that depends on many factors including frequency, amplitude, and viscosity ^13^. *In silico* studies reveal that when exposed to vibrations (0.1-2g), marrow-filled trabecular compartments generate fluid shear stresses up to 2Pa ^13,14^, capable of driving bone cell functions ^15^. Interestingly, while these high magnitude forces are only experienced a few times during the day, bones are bombarded by smaller mechanical signals arising from muscle contractions that generate bone strains ranging between 2 to 10µε ^16^. Exogenous application of small magnitude mechanical regimes in the form of low intensity vibrations (LIV) ranging between 0.1-2g acceleration magnitudes and 20-200Hz frequencies were shown to be effective in improving bone and muscle indices in clinical and preclinical studies ^17^. At the cellular level, our group has reported that application of LIV increases MSC contractility ^18^, activates RhoA signaling ^19^, and results in increased osteogenic differentiation and proliferation of MSCs ^20^.

One of the most actively investigated signaling pathways that regulate the MSC mechanoresponse is the Yes-associated protein (YAP) signaling pathway. YAP depletion in stem cells results in reduced proliferation and osteogenesis ^21,22^. Similarly, depleting YAP from osteoblast progenitors decreases both bone quality and quantity in mice ^23^. Functionally, in response to cytomechanical forces and substrate stiffness, YAP moves from the cytoplasm to the nucleus where it interacts with its co-transcriptional activators such as TEAD to regulate gene expression related to proliferation ^24^. For example, application of substrate strain induces YAP nuclear entry and YAP transcriptional activity which is required to activate proliferation ^25^. While it has been shown that YAP nuclear entry is triggered by soluble factors that increase F-actin contractility such as Lysophosphatidic acid (LPA) ^26,27^, large changes in substrate stiffness ^22^, or substrate stretch ranging from 3% to 15% ^25,28^, it is not known if low magnitude signals like LIV also trigger acute YAP nuclear entry.

Research aimed at studying the effects of microgravity at the cellular level often relies on simulated microgravity (SMG) devices designed to alter the gravitational conditions that cells experience by rotating on one or multiple axes at low speed ^29-31^. SMG decreases MSC proliferation ^32^ and cytoskeletal contractility ^29,33,34^. In this way, application of physical or soluble factors that induce cytoskeletal contractility are commonly used as countermeasures for SMG ^35,36^. SMG also alters nuclear structure. Research from our group has shown that SMG results in reduced levels of integral nuclear proteins such as Lamin A/C and LINC (Linker of Nucleoskeleton and Cytoskeleton) complex elements Sun-2 ^37^. As mechanically induced nuclear shuttling of YAP and its paralog TAZ have been associated with LINC complex function ^38^, SMG also results in decreased nuclear levels of the YAP paralog protein TAZ ^35^. Interestingly, twice daily application of LIV for 20 minutes during SMG recovers both MSC proliferation and levels of nuclear envelope proteins Lamin A/C and Sun-2, suggesting LIV as a potential countermeasure to improve YAP-mediated mechanosignaling in MSCs under SMG.

In this study, using YAP nuclear shuttling as a measure of MSC mechanoresponse, we tested the effect of 72 hours of simulated microgravity (SMG) and daily LIV application (LIV_DT_) on the YAP nuclear entry driven by either acute LIV (LIV_AT_) or Lysophosphohaditic acid (LPA), applied at the end of the 72h period. We hypothesized that SMG-induced impairment of YAP nuclear entry in response to mechanical and soluble factors would be alleviated by daily application of LIV.

## Results

### Acute LIV_AT_ application increases nuclear YAP levels

To quantify the acute YAP nuclear entry in response to LIV, MSCs were plated at density of 5,200 cells/cm^2^ and were allowed to attach for 24hr. Following this, MSCs were subjected to treatment in two groups: control and acute LIV treatment regimen (LIV_AT_). The LIV_AT_ regimen consisted of 5x 20min vibration periods separated by 1hr in between each repetition at room temperature while control samples were treated identically (also taken out of the incubator) but were not vibrated. Immediately after LIV_AT_, samples were immunostained for YAP and DAPI. MATLAB was used to quantify the changes in the nuclear YAP levels. As shown in **Fig.1a**, confocal images showed increased nuclear YAP following the LIV_AT_ treatment. Analysis of confocal images to quantify nuclear YAP intensity shown in **Fig.1b** revealed a 32% increase in the nuclear YAP levels in the LIV_AT_ samples as compared to the control samples (p<0.0001). We also used C2C12 myoblasts to confirm the LIV_AT_ induced YAP nuclear entry on a second cell line, quantitative analysis of confocal images showed a 40% increase of nuclear YAP in LIV_AT_ samples compared to controls (**Fig.S1**). As both LIV-induced focal adhesion signaling, initiated by focal adhesion kinase (FAK) phosphorylation at Tyr 397 residue ^19^, and YAP nuclear entry in response to substrate strain ^28^ requires intact LINC function, disabling LINC function via a dominant negative overexpression of Nesprin KASH (Klarsicht, ANC-1, Syne homology) fragment both decreased basal nuclear YAP levels by 34% (p<0.0001) and impeded the LIV-induced YAP nuclear entry when compared to empty plasmid (**Fig.S2**). FAK phosphorylation at Tyr 397 residue (pFAK) was blocked via a FAK inhibitor (FAKi) PF573228 (3µM) 1hr prior to LIV_AT_ treatment as previously described ^19^ and stained against DAPI and YAP (**Fig.S3a**). FAKi inhibited the LIV_AT_ induced pFAK and decreased its basal levels (**Fig.S3b**). As shown in **Fig.S3c**, measuring nuclear YAP levels showed that LIV_AT_ induced YAP nuclear entry was not affected by FAKi when compared DMSO treated controls.

**Figure 1.**
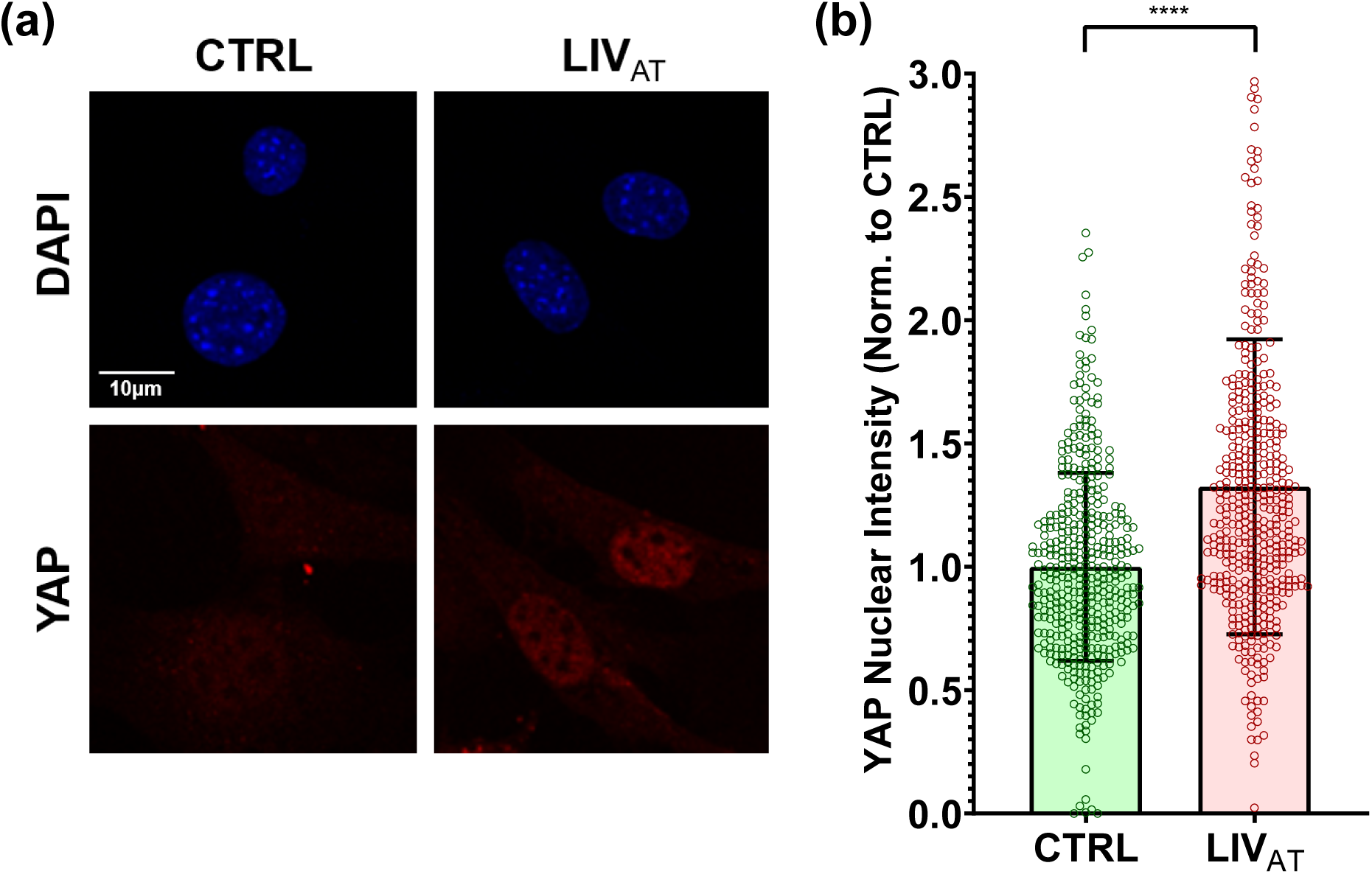
Acute LIV_AT_ application increases nuclear YAP levels. (a) MSCs were subjected to LIV_AT_ and stained with DAPI (blue) and YAP (red). Confocal images indicated an increased nuclear YAP levels following acute LIV_AT_ applied as five 20min vibration periods separated by 1hr. (b) Quantitative analysis of confocal images showed a 32% of increase of nuclear YAP in LIV_AT_ samples compared to controls. n>400/grp, group comparison was made a Mann-Whitney U-test, *p<0.05, **p<0.01, ***p<0.01, ****p<0.0001.

### Basal nuclear YAP levels decreased by SMG were rescued by daily application of LIV_DT_

We next tested whether SMG decreases basal YAP levels and whether a daily LIV treatment regimen (LIV_DT_), applied in parallel with SMG, could alleviate decreased YAP in the nucleus. As we reported previously, LIV_DT_ consisting of 2x 20min vibrations applied every 24 hours during the 72h period of SMG. This LIV_DT_ regimen was effective at restoring MSC proliferation and whole cell YAP levels when applied in conjunction with SMG ^37^. MSCs were plated at a density of 1,700 cells/cm^2^ in 9cm^2^ tissue culture SlideFlasks (Nunc, #170920) and were allowed to attach for 24h, after which point, the flasks were filled completely with growth medium, sealed, and subjected to 72h of treatment followed by immunostaining for YAP and nuclear staining using DAPI. During the 72h treatment period, MSCs were divided into three groups: control samples, SMG samples which were subjected to the 72h SMG alone, and SMG+LIV_DT_ samples which were subjected to both the 72h SMG regimen and the daily LIV_DT_ regimen. Representative images for YAP and DAPI stained images are shown in **Fig.2a.** As depicted in **Fig.2b**, quantitative analysis of confocal images revealed a 42% decrease in the nuclear YAP intensity of the SMG group as compared to non-SMG controls (p<0.0001). Compared to the SMG group, the LIV_DT_ group increased nuclear YAP levels by 67% (p<0.0001) and there was no significant difference between nuclear YAP levels of LIV_DT_ treated MSCs and non-SMG controls.

**Figure 2.**
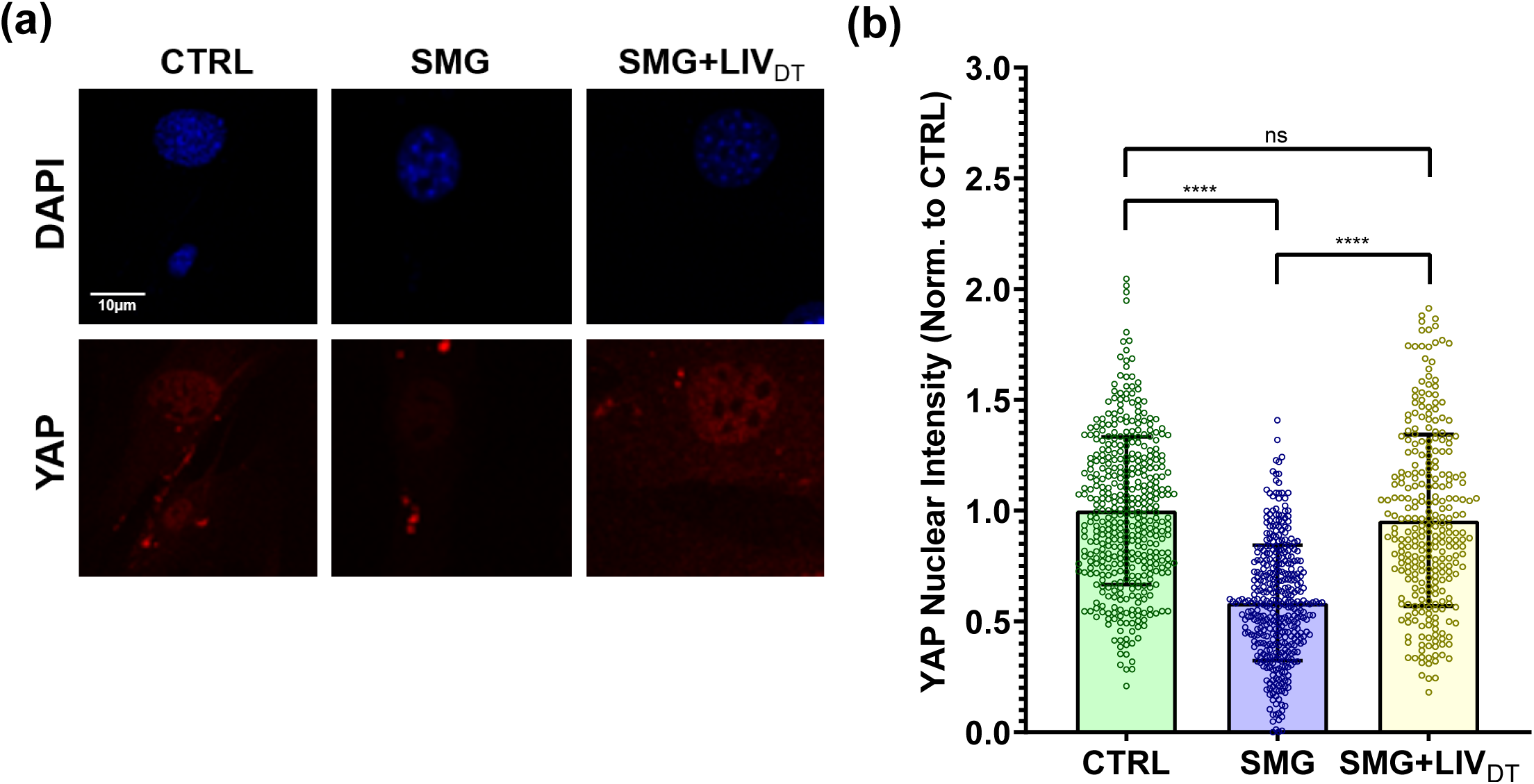
Basal nuclear YAP levels decreased by SMG were rescued by LIV_DT_. (a) MSCs were subjected to SMG, and SMG+LIV_DT_ over 72h period and stained with DAPI (blue) and YAP (red). (b) Quantitative analysis showed a 42% decrease of nuclear YAP levels in the SMG group compared to control levels. The SMG+LIV_DT_ group showed a 67% increase of nuclear YAP when compared to the SMG group. There was no statistically significant difference between CTRL and SMG+LIV_DT_ groups. n>100/grp. Group comparisons were made via Kruskal-Wallis test followed by Tukey multiple comparison, *p<0.05, **p<0.01, ***p<0.01, ****p<0.0001.

### LIV_AT_-induced YAP nuclear entry decreased by SMG was not restored by daily LIV_DT_ application

As SMG decreased basal nuclear YAP levels, we next tested whether SMG decreases LIV_AT_-induced YAP mechanosignaling (i.e. nuclear shuttling). Since LIV_DT_ was able to restore nuclear YAP levels (**Fig.2**), the SMG+LIV_DT_ group was added to evaluate the effect of LIV_DT_ on the SMG response. A schematic of the experimental design is given in **Fig.3**. MSCs were divided into six groups in which the CTRL, SMG, SMG+LIV_DT_ groups were treated with ±LIV_AT_ at the end of 72h and nuclear YAP levels were measured. As shown in **Fig.4**, SMG alone decreased basal nuclear YAP levels by 37% (p<0.0001) which were restored back to control levels in the SMG+LIV_DT_ group. As depicted in **Fig.4**, +LIV_AT_ increased nuclear YAP levels in the CTRL, SMG and SMG+LIV_DT_ groups by 50%, 69% and 22%, respectively (p<0.0001) while exhibiting the smallest increase in the SMG+LIV_DT_. As a result, final nuclear YAP levels in the SMG+LIV_DT_+LIV_AT_ group was not significantly different from the SMG+LIV_AT_ and 23% lower than the LIV_AT_ group (p<0.0001). Representative confocal images are presented in **Fig.S4**.

**Figure 3.**
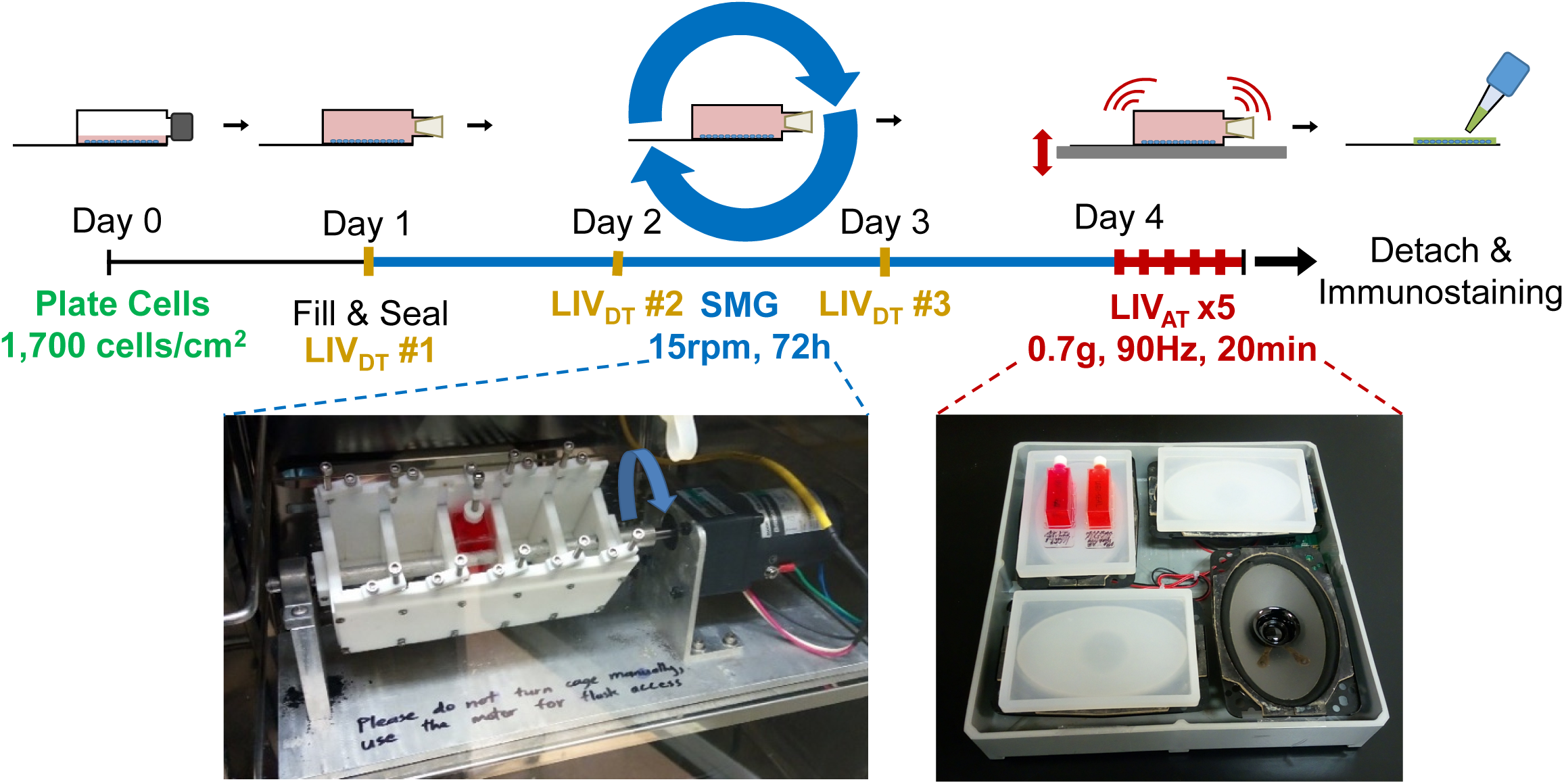
Experimental design of combined SMG, LIV_AT_ and LIV_DT_ application. MSCs were subcultured and plated in SlideFlasks and allowed to attach for 24h before SlideFlasks were filled with culture medium and sealed. Treatment regimen for MSC’s involved 72h of SMG (blue). LIV_DT_ regimen consisted of one treatment cycle every 24hr during SMG treatment with each cycle consisting of 2x 20min LIV with an hour in between (yellow). LIV_AT_ regimen was applied after 72h SMG treatment period and consisted of 5x 20min LIV with an hour in between each (red). For LIV application, MSCs plated in SlideFlasks were placed in LIV device constructed in the lab previous to this research. Vibrations were applied at peak magnitudes of 0.7 g at 90 Hz at room temperature. Control samples were treated the same but were not vibrated. For SMG application, MSCs plated in SlideFlasks were secured in lab custom-built clinostat inside incubator. The clinostat subjected the MSCs to constant 15 RPM rotation simulated microgravity. After treatment, flasks were removed for immunofluorescence staining.

**Figure 4.**
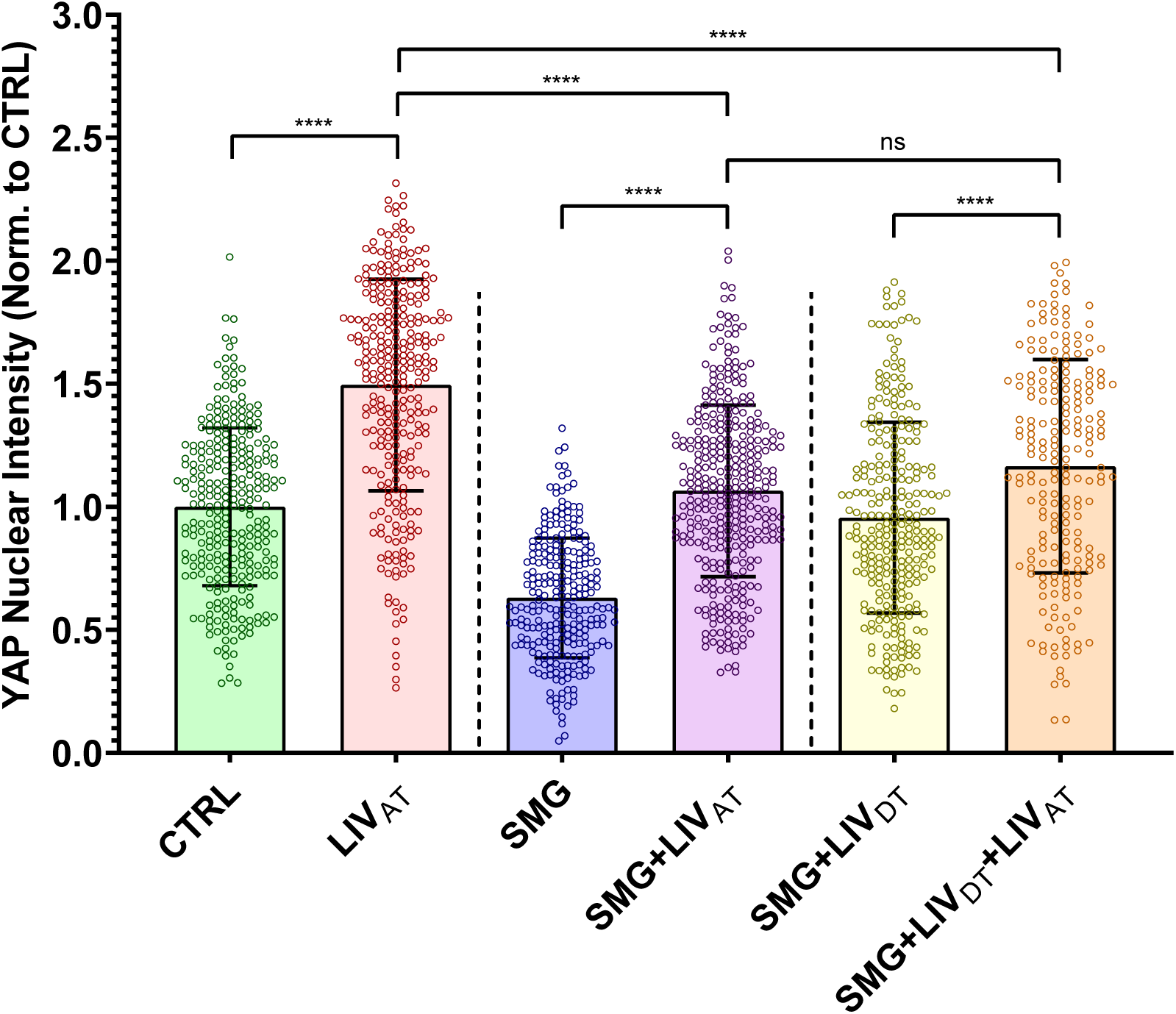
LIV_AT_-induced YAP nuclear entry decreased by SMG was not restored by daily LIV_DT_ application. MSCs were subjected to either CTRL, SMG, SMG+LIV_DT_ over 72h period were subsequently treated with LIV_AT_. Quantitative analysis of confocal images showed that SMG alone decreased basal nuclear YAP levels by 37% which were increased back to control levels in the SMG+LIV_DT_ group. +LIV_AT_ increased nuclear YAP levels in the CTRL, SMG and SMG+LIV_DT_ groups by 50%, 69% and 22%, respectively. Nuclear YAP intensity in the SMG+LIV_DT_+LIV_AT_ group remained not significantly different from the SMG+LIV_AT_ and 23% smaller than the LIV_AT_ group. n>200/grp. Group comparisons were made via Kruskal-Wallis test followed by Tukey multiple comparison, *p<0.05, **p<0.01, ***p<0.01, ****p<0.0001.

### LPA treatment increases nuclear YAP levels

As SMG+LIV_DT_ treatment did not improve the acute LIV_AT_ response when compared to the SMG group, we next considered a soluble regulator of cytoskeletal contractility, LPA ^20^. To test the effect of LPA on the acute YAP nuclear entry, two LPA concentrations (50µM and 100µM) were compared against control samples. Shown in **Fig.5**, nuclear YAP levels were almost doubled under a two-hour exposure to 50µM LPA and 100µM LPA treatments with 99% and 107% increases as compared to the control samples (p<0.0001). Nuclear YAP levels for 50µM LPA and 100µM LPA treatments were not significantly different. Therefore, we chose to use 50µM LPA treatment in the subsequent experiments.

**Figure 5.**
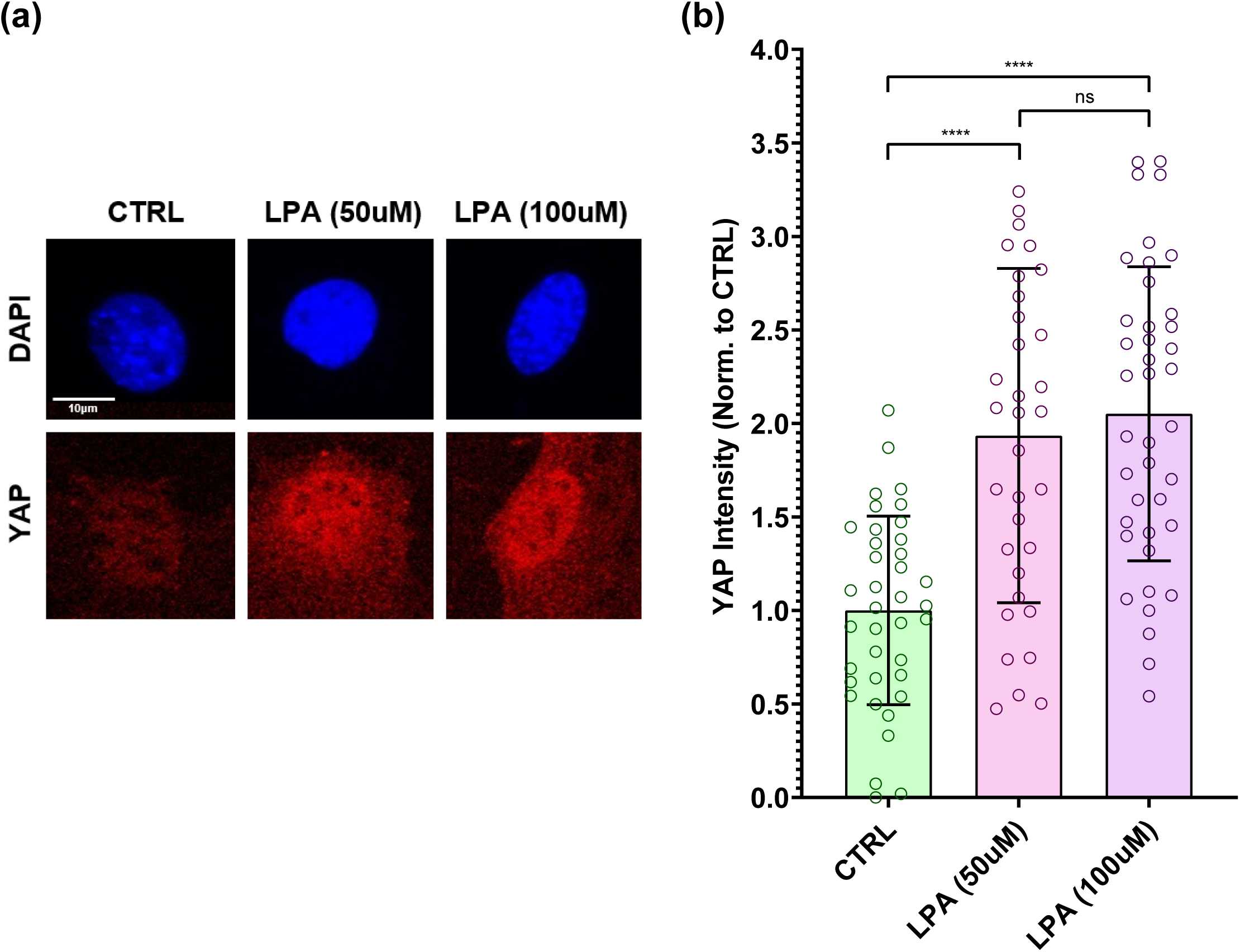
LPA treatment increases nuclear YAP levels. (a) Representative confocal images of DAPI (blue) and YAP (red) stained MSCs with or without LPA treatment. MSCs were subjected to LPA addition at 50µM and 100µM concentrations. (b) Quantitative analysis of confocal images revealed a 99% and a 107% increase in the 50µM LPA and 100µM LPA treatments compared to DMSO treated controls, respectively. Nuclear YAP levels for 50µM LPA and 100µM LPA treatments were not significantly different. n>30/grp. Group comparisons were made via Kruskal-Wallis test followed by Tukey multiple comparison, *p<0.05, **p<0.01, ***p<0.01, ****p<0.0001.

### LPA-induced YAP nuclear entry decreased by SMG was alleviated by daily LIV_DT_ application

In order to evaluate whether LIV_DT_ can restore LPA-induced YAP nuclear entry after SMG, 50µM LPA dissolved in DMSO or DMSO as vehicle control were added to the samples at the end of the 72h treatment of either CTRL, SMG or SMG+LIV_DT_ treatments. The CTRL group, SMG group, and SMG+LIV_DT_ group were subjected to the same treatment as in the previous experiments and displayed similar results. As depicted in **Fig.6**, +LPA increased nuclear YAP levels in the CTRL, SMG and SMG+LIV_DT_ groups by 105%, 67% and 43% respectively (p<0.0001). While final YAP nuclear levels in SMG+LIV_DT_+LPA remained 70% higher than SMG+LPA group (P<0.0001), it remained 29% lower than the LPA group (P<0.0001).

**Figure 6.**
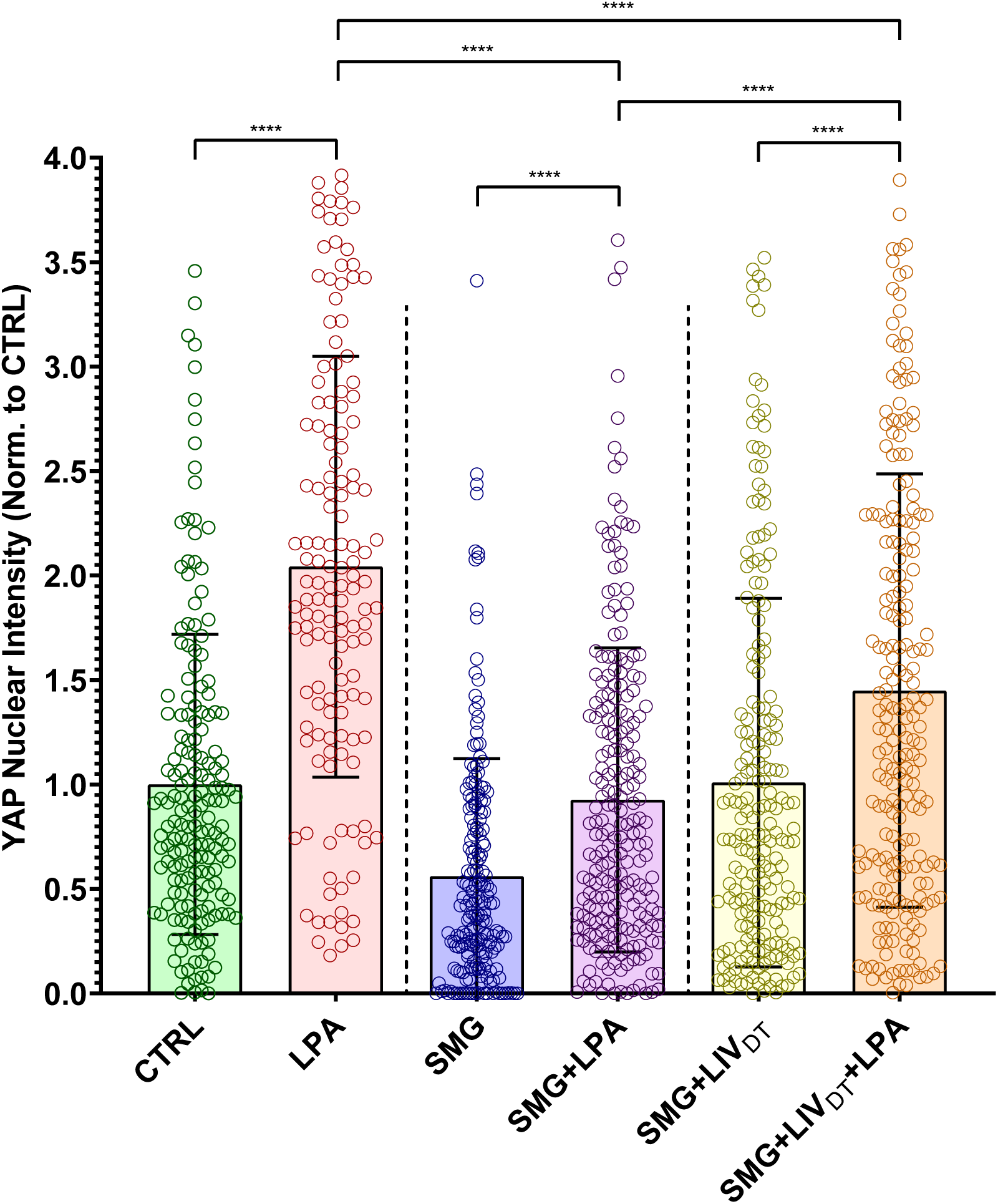
LPA-induced YAP nuclear entry decreased by SMG was alleviated by daily LIV_DT_ application. MSCs were subjected to SMG, and parallel SMG+LIV_DT_ over 72h period at the end of 72h, samples were treated with either LPA (50µM) or DMSO. Quantitative analysis of confocal images revealed that LPA addition increased nuclear YAP levels by 105%, 67% and 43% in the CTRL, SMG and SMG+LIV_DT_ when compared to DMSO controls. When compared to nuclear YAP intensity of the LPA treatment alone, SMG+LPA and SMG+LIV_DT_+LPA samples were 55% and 29% lower, respectively. YAP nuclear levels in SMG+LIV_DT_+LPA remained 70% higher than SMG+LPA group. n>100/grp. Group comparisons were made via Kruskal-Wallis test followed by Tukey multiple comparison, *p<0.05, **p<0.01, ***p<0.01, ****p<0.0001.

### MSC stiffness and structure remain intact under SMG and SMG+LIV_DT_ treatments

As YAP mechanosignaling of SMG+LIV_DT_ MSCs remained below control levels in response to both LIV_AT_ and LPA, we quantified the effects of SMG and SMG+LIV_DT_ on the cell stiffness, F-actin intensity, cell area and nuclear area. AFM testing was used to quantify the elastic modulus of the nucleus by measuring load-displacement curves on top of the nucleus. AFM tests shown in **Fig.7a** indicated a 21% and 27% stiffness decrease in the SMG or SMG+LIV_DT_ groups but differences were not significant. Quantified from confocal images (**Fig.7b**), mean F-actin intensities for all the cells in each imaging field were quantified by dividing the mean F-actin intensity to the number of nuclei in each imaging field. Shown in **Fig.7c**, SMG and SMG+LIVDT treated MSCs revealed 36% and 30% decreases in the mean F-actin intensity per cell, respectively but the differences were not statistically significant. We have further quantified nuclear area as a measure of cyto-mechanical forces on the nucleus ^39^. Shown in **Fig.7d**, analysis of cross-sectional area of cell nuclei using DAPI stained images revealed no significant effect on average nuclear size by either SMG or combined SMG+LIV_DT_ treatment compared to control levels.

**Figure 7.**
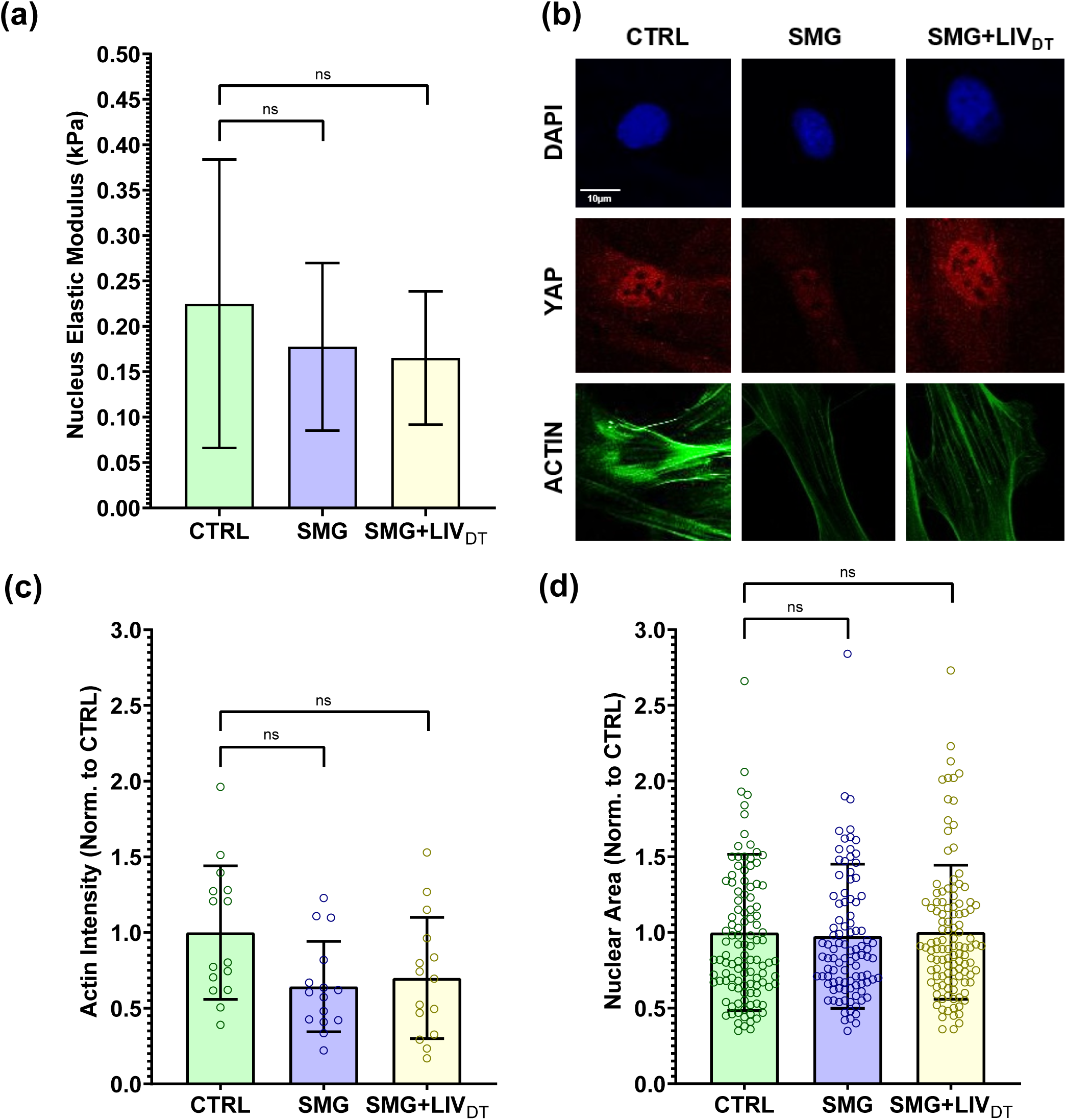
MSC stiffness and structure remain intact under SMG and SMG+LIV_DT_ treatments. MSCs were subjected to SMG and parallel SMG+LIV_DT_ over a 72h period. (a) Compared to CTRL samples, AFM measurement of the elastic moduli of SMG and SMG+LIV_DT_ treated MSCs revealed apparent decreases in elastic modules that were 21% and 27% below control levels, measured differences were not statistically significant. n=10/grp. (b) Quantification of confocal images show that, (c) mean F-actin intensity of SMG and SMG+LIV_DT_ treated MSCs revealed decrease of 36% and 30% below control levels, measured differences were not statistically significant. n=15/grp. (d) No significant effects of either SMG or LIV_DT_ treatment on the average nucleus size were found. n>100/grp. Group comparisons were made via Kruskal-Wallis test followed by Tukey multiple comparison, *p<0.05, **p<0.01, ***p<0.01, ****p<0.0001.

## Discussion

The mechanical forces that the bone and muscle cells are subjected to on Earth and in microgravity are complex and remain incompletely understood. At the same time, it is clear that these forces are required for healthy tissue growth and function. The complexity of these forces makes it difficult to design experiments that comprehensively simulate *in vivo* conditions. While the *in vitro* experiments utilizing SMG and LIV treatments used in this study are limited in this way and do not entirely correlate with the physiological behavior of these cells *in vivo*, the experiments presented here remain useful for testing cell behavior under well-defined conditions.

In this study, we focused on YAP mechanosignaling of MSCs. The first experiments demonstrated that repeated LIV_AT_ application over six hours was capable of stimulating YAP entry into the nucleus in both MSCs (**Fig.1**) and in C2C12 cell line (**Fig.S1**). These findings suggest that similar to high magnitude substrate strains ^25^ smaller mechanical signals such as LIV can be effective at increasing nuclear YAP levels. Further, in agreement with earlier reports utilizing uniaxial strain ^28^, LIV_AT_-induced increase in nuclear YAP levels also required functional LINC complexes (**Fig.S2**). Integrin related FAK signaling have been shown to promote YAP nuclear levels in the proliferative descendants of stem cells and that FAK inhibitor PF573228 decreased nuclear YAP in these cells ^40^. Similarly, inhibiting integrin engagement via blocking FAK phosphorylation in Tyr 397 residue via FAKi also mutes the increase of GTP-bound RhoA levels in LIV treated MSCs ^19^. While we confirmed the loss of phosphorylation in Tyr 397 at both basal level and in response to LIV_AT_ (**Fig.S3b**), FAKi treatment changed neither basal levels nor the LIV_AT_-induced increase in nuclear YAP (**Fig.S3c**), suggesting a FAK independent mechanism. In these experiments, we did not compare LIV_AT_ with strain because the application of 5 to 15% stretch onto sealed culture flasks was not technically possible without significantly altering experimental conditions. Instead, LPA addition served as the best option for applying a simple mechanical stimulation in order to evaluate the YAP mechanotransduction. LPA is a phospholipid derivative signaling molecule which is capable of causing the simulation of static transient stretch of a cell by increasing the contractility of the cytoskeleton ^20,41^. The first experiments with LPA served to verify that the simulation of stretch via increased cytoskeleton contractility was capable of triggering YAP entry into the nucleus and the analysis methods utilized here were capable of detecting this response (**Fig.5**).

The first SMG experiments confirmed a clear decrease of basal nuclear YAP levels. Interestingly, SMG treated cells remained responsive, as both LIV_AT_ and LPA treatments were able to increase the nuclear YAP levels at the end of acute stimulation period (< 6h). However, final nuclear YAP levels in SMG treated MSCs remained significantly lower when compared to non-SMG groups (**Fig.2, 4 & 6**). These findings suggested that the YAP mechanosignaling apparatus of MSCs, to some extent, was intact under SMG. When applied in parallel to SMG, daily LIV_DT_ treatment was able to restore basal YAP levels in the cell nucleus (**Fig.2, 4 & 6**) measured 24h after the final LIV_DT_ treatment. This increase in nuclear levels supported our earlier report that showed sustained recovery of MSC proliferation by LIV_DT_ ^37^.

Interestingly, this increase in basal nuclear YAP levels under LIV_DT_ was accompanied by a reduced MSC response to LIV_AT_ treatment (**Fig.4**). When SMG+LIV_DT_ treated MSCs were subjected to LIV_AT_, the increase in nuclear YAP from the non-LIV_AT_ control was only 22% (p<0.0001), which was small compared to the 77% increase seen in the SMG groups in response to LIV_AT_. As a result of this smaller increase in the SMG+LIV_DT_ group, there was no measurable difference between SMG and SMG+LIV_DT_, samples that were subjected to LIV_AT_. It has been previously reported that an application of multiple LIV bouts separated by a refractory period is more effective at activating mechano-signaling pathways such as βcatenin ^42^. It is possible that long term application of LIV_DT_ results in cell structural adaptations that serve to reduce MSC responsiveness to LIV_AT_ treatment. To test this possibility, we replaced LIV_AT_ with an LPA treatment. When LIV_AT_ was replaced by LPA treatment (**Fig.6**), responsiveness of SMG+LIV_DT_ treated MSCs almost doubled to 43% (compared to 22% in response to LIV_AT_) and was significantly higher than the SMG+LPA group (p<0.0001), suggesting that LIV_DT_ increases the YAP-mechanosignaling in response to LPA.

Absolute nuclear YAP intensity in the SMG+LIV_DT_+LIV_AT_ group, however, remained below the LIV_AT_ group (p<0.0001). Previously published findings using the same treatment protocols suggested that the total cellular YAP levels decreased by SMG were restored to control levels by daily LIV ^37^. This indicates that total availability of YAP protein was not responsible for this difference between the SMG+LIV_DT_+LIV_AT_ and the LIV_AT_ groups. In regards to other potential effects of SMG on the components of the mechanosignaling mechanism, one current prevailing hypothesis suggests a role for nuclear pore opening in response to cyto-mechanical forces ^38^ which may be affected by changes in the nuclear stiffness. To test this possibility, we performed additional AFM and imaging experiments. While the AFM measured nuclear stiffness was 24% lower in the SMG and SMG+LIV_DT_ groups on average, we were unable to identify any statistically significant effects of SMG or LIV_DT_ treatment on nuclear stiffness. There was also slight F-actin intensity decreases in both the SMG and SMG+LIV_DT_ groups which were also not significant (**Fig.7**). Similarly, cell and nuclear area were not affected. While our results were not able to detect any changes in nuclear stiffness, considering the significant role that the nuclear membrane plays as a mechanical structural component in the cell’s interpretation of mechanical stimulus ^43,44^, more detailed future studies are needed to study the effects of SMG on the nuclear envelope and nuclear structure.

In summary, while the restoration of basal nuclear levels and improvement of LPA induced YAP nuclear entry under daily LIV_DT_ treatment identify LIV as a possible countermeasure to improve MSC response under the detrimental effects of simulated microgravity, future studies are required to understand why acute YAP nuclear entry in response to mechanical and soluble factors remain less responsive.

## Methods and Materials

### Cell Culture

Primary mouse bone marrow derived MSC’s were extracted as previously described ^37,45^. C2C12 mouse myoblasts were derived from muscle satellite cells. MSCs were subcultured and plated in Iscove modified Dulbecco’s cell culture medium (IMDM, 12440053, Gibco) with 10% fetal calf serum (FCS, S11950H, Atlanta Biologicals) and 1% pen/strep. C2C12s were subcultured and plated in Dulbecco’s modified Eagle’s medium (DMEM, DML09, Caisson Laboratories) with 10% fetal calf serum (FCS, S11950H, Atlanta Biologicals) and 1% pen/strep. MSCs were subcultured in 9cm^2^ culture dishes at a density of 5,200cells/cm^2^ for 1day experiments and 1700cells/cm^2^ for 3day experiments, while C2C12s were plated at a density of 10,000cells/cm^2^. Experimental cells were plated and given 24h to attach to the mounting surface prior to experiments. Cell passages for both MSCs and C2C12s used for experiments were limited to P7-P15.

### Low Intensity Vibrations Treatment

SlideFlasks with plated MSCs were filled completely with culture medium and placed in LIV device designed and used in previous research (**Fig.3**) ^37^. LIV device subjected cells to low intensity 90 Hz lateral vibrations at 0.7g at room temperature. MSCs were vibrated for 20min intervals separated over time. LIV_AT_ regimen was applied after 72h treatment period and consisted of 5x 20min LIV with an hour in between each. Daily LIV_DT_ regimen consisted of 3x treatments in parallel with SMG treatment each consisting of 2x 20min LIV with 2h in between.

### Simulated Microgravity Treatment

SlideFlasks (Nunc, #170920) with plated MSCs were filled completely with culture medium (**Fig.3**) and placed in a clinostat SMG device. The clinostat shown is a redesign of a custom-made clinostat described in previous research ^37^ with a new flask holder casing capable of holding SlideFlasks and is also autoclavable. The clinostat was used to subject the MSCs to constant 15 RPM SMG for 72h.

### Immunofluorescence Staining and Image Analysis

Immediately after mechanical treatment, MSCs plated in Slideflasks were removed from treatment, and the SlideFlasks were disassembled in order to stain the MSCs on the slides (**Fig.3**). The MSCs were fixed with 4% paraformaldehyde, then washed and permeabilized with 0.05% Triton X-100 in PBS, followed by immunostaining with YAP specific antibody (YAP (D8H1X) Rabbit mAb, Cell Signaling Technologies) and Alexa Fluor red secondary antibodies (Donkey anti-Rabbit IgG (H+L) Cross-Adsorbed Secondary Antibody, Alexa Fluor Plus 594 for all experiments prior to usage of LPA. Subsequently, Donkey anti-Rabbit IgG (H+L) Cross-Adsorbed Secondary Antibody, Alexa Fluor Plus 633 was used). Nuclear DNA was labeled via DAPI (Vectashield Mounting Medium, Vector Laboratories). Stained samples were imaged with a Leica TCS SP8 confocal microscope (40x, HC PL APO CS2 Oil Immersion) prior to usage of LPA, after this Zeiss LSM 510 Meta Confocal Microscope (40x, HC PL APO CS2 Oil Immersion). Exported images were used to quantify relative YAP levels within each nuclei (nuclear regions traced by DAPI stained nucleus) via custom-made MATLAB program (The MathWorks, Natick, MA). DAPI images were analyzed using an edge-detection algorithm in order to determine the nuclear area for each cell. The nuclear outline was then used as a mask to quantify the average pixel intensity of the YAP stain within the nuclei of each individual cell. (n=50-100 nuclei/sample).

### Atomic Force Microscopy

Bruker Dimension FastScan AFM was used for collection of the atomic force measurements. Tipless MLCT-D probes with a 0.03 N/m spring constant were functionalized with 10 µm diameter borosilicate glass beads for force collection. The AFM’s optical microscope was used to locate individual live MSCs plated on the SlideFlask slides with the flask section removed for access to the cells. The nucleus of each cell was tested with at least 3 seconds of rest between each test. In each test, three force-displacement curves were obtained (ramping rate: 2 µm/sec over 2 µm total travel, 1 µm approach, 1 µm retract), which were analyzed using Nanoscope software with the implementation of a best-fit curve to a Hertzian (spherical) model (optimized such that R^2 value was greater than 0.95, or p<0.05) to obtain elastic moduli of nuclear membrane of individual nuclei.

### Western Blotting

Western blotting was performed as previously described.^23,26,27,64^ 20μg of lysed cell protein from each sample was run on a 10% polyacrylamide gels, transferred onto a polyvinylidene difluoride (PVDF) membranes, blocked with 5% (w/v) milk for 1h. After washing, primary antibodies were incubated overnight at 4°C with. Protein bands were visualized via horseradish peroxidase-conjugated secondary antibodies (1: 5,000, Cell Signaling) and ECL plus chemiluminescence kit (Amersham Biosciences, Piscataway, NJ) and scanned using C-DiGit blot scanner (Licor, Lincoln, NE). All blots derive from the same experiment and were processed in parallel.

### Statistical analysis

All data analysis results were displayed graphically based on the mean value with standard error bars. Differences between treatments were not assumed to follow a Gaussian distribution. Therefore, group differences were identified via either non-parametric two-tailed Mann-Whitney U-test (**Fig.1a**) or Kruskal-Wallis test followed by Tukey multiple comparison (**Fig.2b, 4, 5, 6, 7, &S1, S2, S3**). P-values of less than 0.05 were considered significant.

## Data availability

The datasets generated and/or analyzed during the current study are available from the corresponding author on reasonable request.

## Ethics

All methods were carried out in accordance with relevant guidelines and regulations of Boise Institutional Animal Care and Use Committee and Institutional Biosafety Committee. All procedures were approved by Boise State University Institutional Animal Care and Use Committee, and Institutional Biosafety Committee.

## Acknowledgements

This study was supported by NASA ISGC NNX15AI04H, NIH R01AG059923, and 5P2CHD086843-03, P20GM109095, P20GM103408 and NSF 1929188.

## Contributions

**Thompson, M** experimental methods, data analysis/interpretation, manuscript writing, final approval of manuscript.

**Woods, K** data analysis/interpretation, final approval of manuscript.

**Newberg, J** experimental methods, final approval of manuscript

**Oxford JT**, financial support, final approval of manuscript

**Uzer, G** concept/design, financial support, data analysis/interpretation, manuscript writing, final approval of manuscript

## Supplementary Information

**Figure S1.**
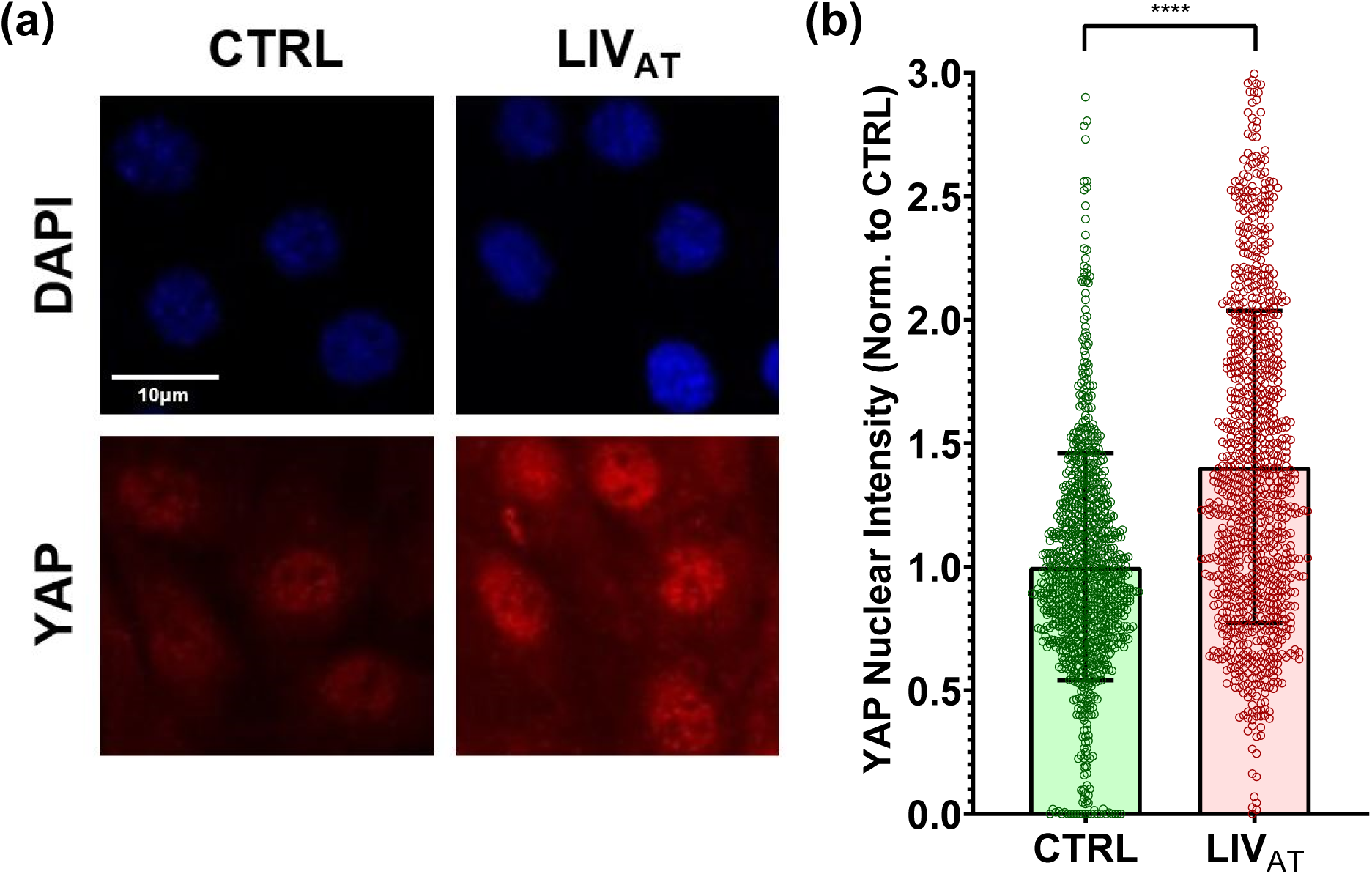
LIV_AT_ treatment increases nuclear YAP in C2C12 cells. (a) C2C12 cells were subjected to LIV_AT_ and stained with DAPI (blue) and YAP (red). Confocal images displayed increased nuclear YAP levels following LIV_AT_ treatment. (b) Quantitative analysis of confocal images showed a 40% increase of nuclear YAP in LIV_AT_ samples compared to controls. n>900/grp, group comparison was made using a Mann-Whitney U-test, *p<0.05, **p<0.01, ***p<0.01, ****p<0.0001.

**Figure S2.**
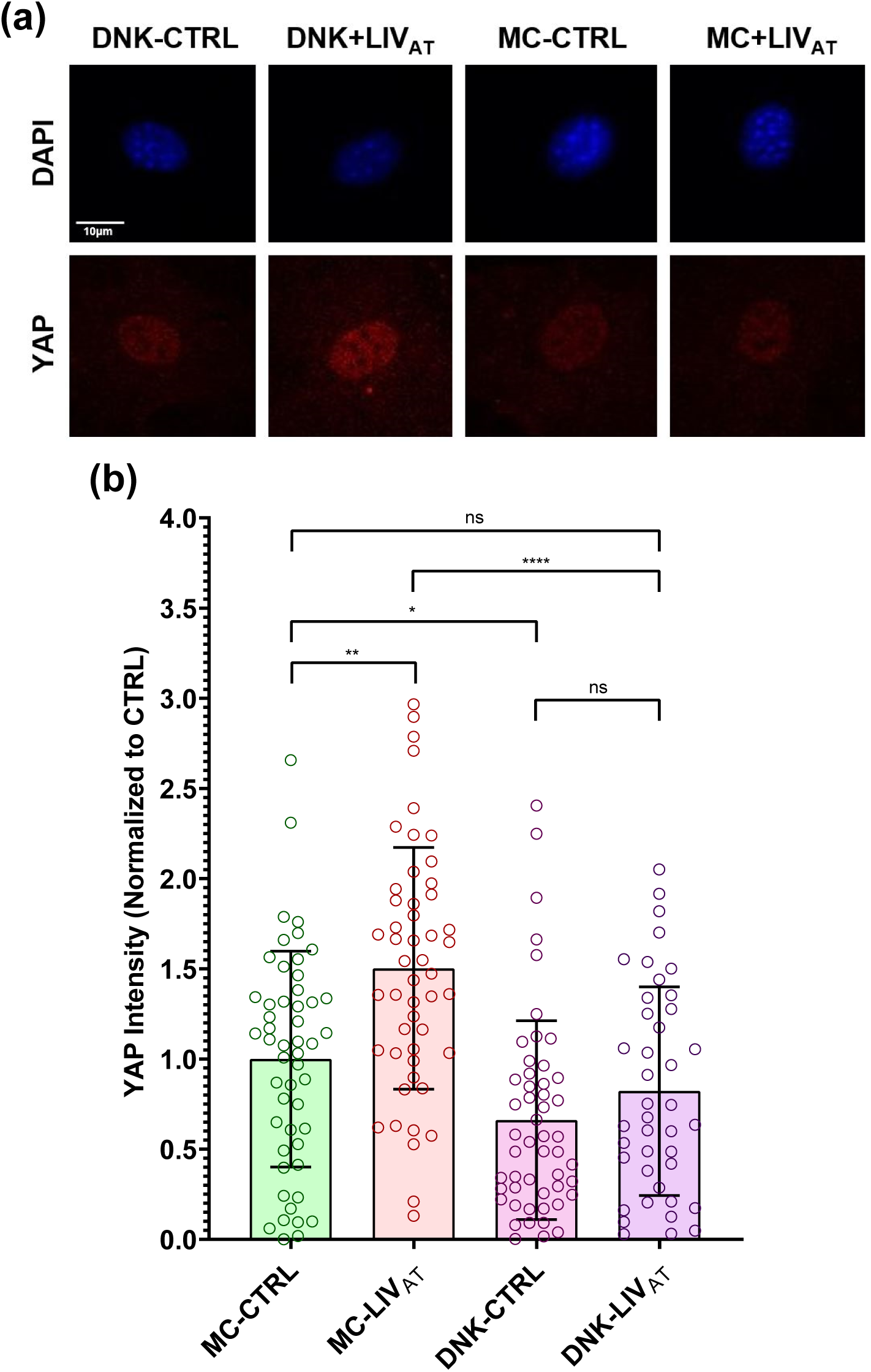
LINC complex disruption decreases nuclear YAP levels and reduces LIV_AT_-induced YAP nuclear entry. (a) Plasmids harboring either a dominant negative KASH domain of Nesprin (DNK) to disable LINC complex function or empty mCherry control (MC) were overexpressed in MSCs. Following puromycin selection, MC or DNK expressing MSCs were subjected to LIV_AT_ and stained against DAPI (blue) and YAP (red). (b) Quantitative analysis of confocal images revealed a 49% increase of nuclear YAP following LIV_AT_ in MC expressing control MSCs. Basal YAP levels of the DNK-CTRL group were 34% lower compared to MC-CTRL and LIV_AT_ treatment failed to significantly increase nuclear YAP over DNK-CTRL (18%, NS). n>30/grp, group comparisons were made via Kruskal-Wallis test followed by Tukey multiple comparison, *p<0.05, **p<0.01, ***p<0.01, ****p<0.0001.

**Figure S3.**
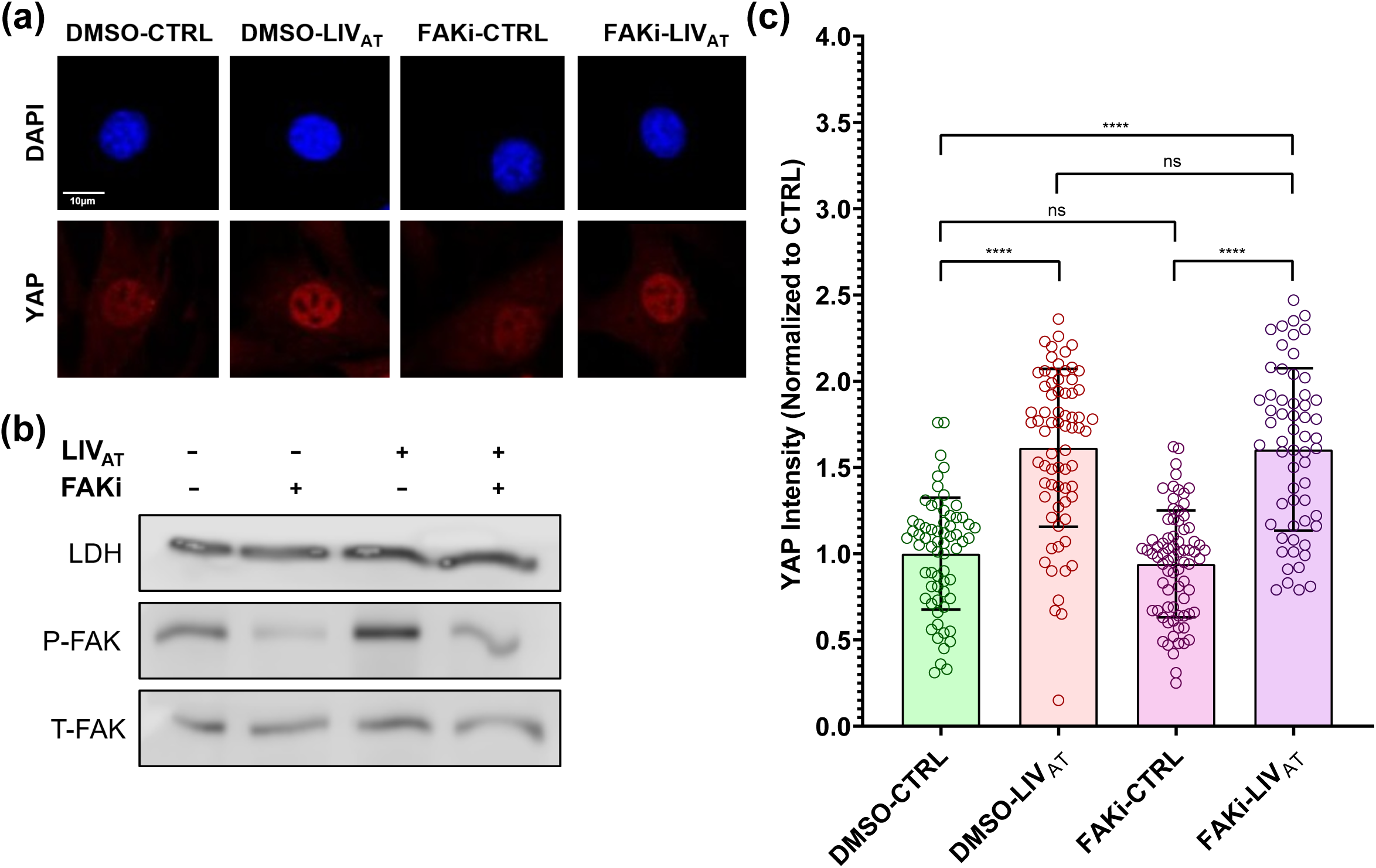
Blocking FAK phosphorylation at Tyr 397 does not limit LIV_AT_ induced YAP nuclear entry. Dimethyl sulphoxide (DMSO) or Tyr 397 specific FAK inhibitor (FAKi) PF573228 (3µM) was added to MSCs in culture medium for 1h prior to LIV_AT_ or control treatments. (a) Confocal images of YAP showed more intense nuclear YAP staining of LIV_AT_ treated MSCs but no apparent effect of FAKi when compared to DMSO (b) FAKi application 1hr prior to LIV_AT_ treatment inhibited the LIV_AT_ induced FAK phosphorylation at Tyr 397 and decreased the basal levels (c) Quantitative analysis of confocal images revealed a 61% increase of nuclear YAP in both the DMSO-LIV group and a 60% increase in the FAKi-LIV_AT_ group compared to the DMSO-CTRL group. Differences between the DMSO-CTRL and FAKi-CTRL groups and between the DMSO-LIV and FAKi-LIV_AT_ groups were not significant. n>50/grp. Group comparisons were made via Kruskal-Wallis test followed by Tukey multiple comparison, *p<0.05, **p<0.01, ***p<0.01, ****p<0.0001.

**Figure S4.**
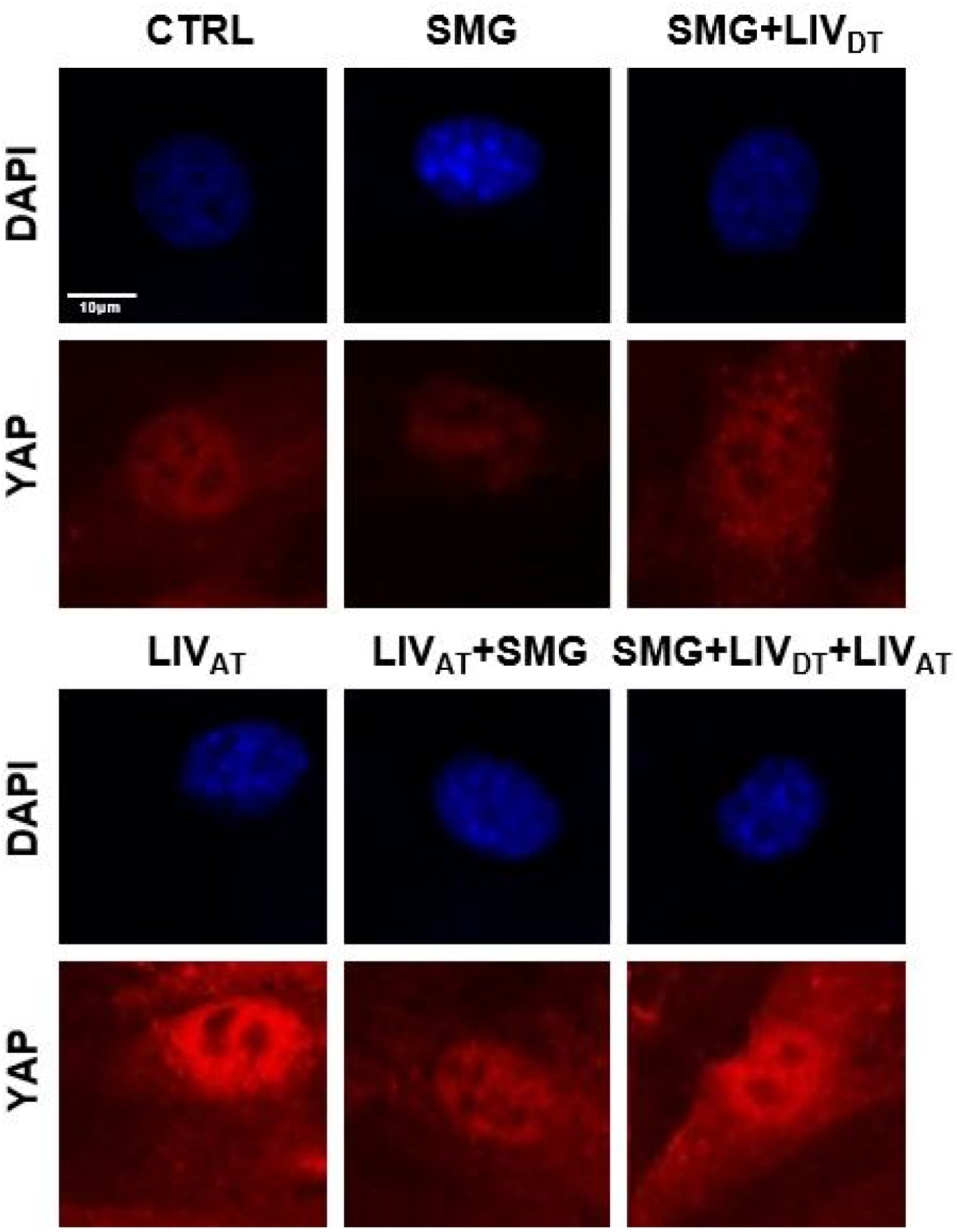
Confocal images for SMG/LIV_DT_/LIV_AT_ treatments in Figure 4.

**Figure S5.**
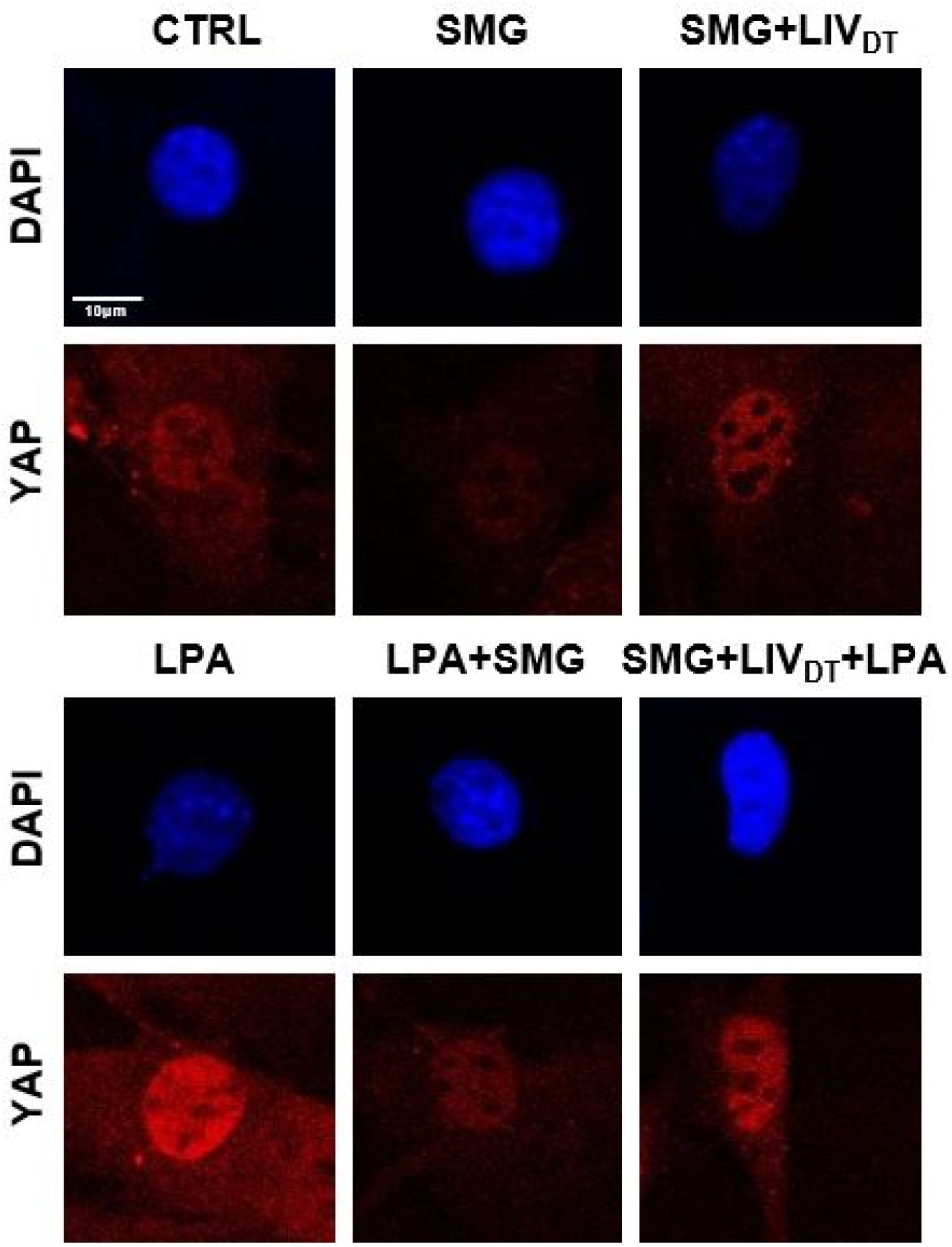
Confocal images for SMG/LIV_DT_/LPA treatment in Figure 6.

**Table S1:**
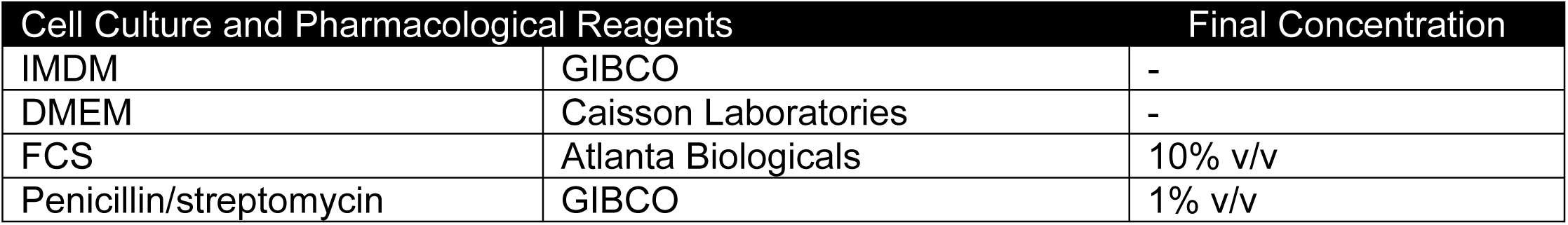
Cell culture and pharmacological reagents and their final concentrations.

**Table S2:**
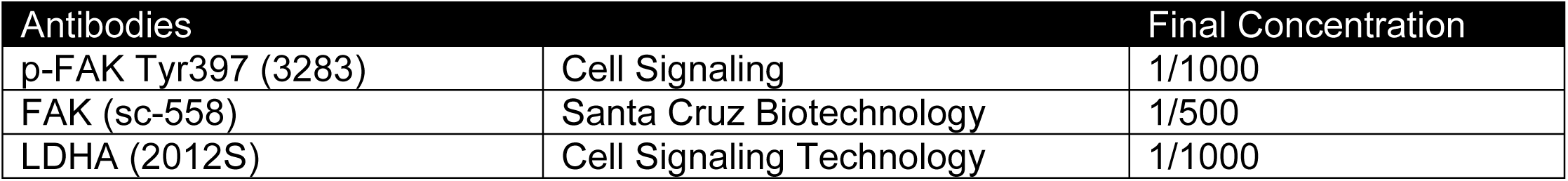
Antibodies used and their final concentrations for western blots.

**Table S3:**
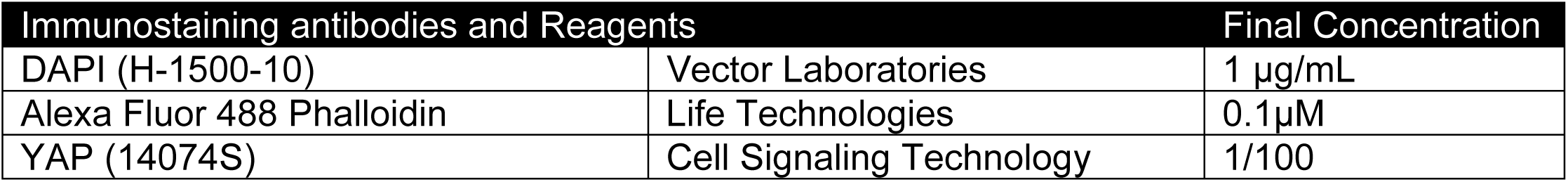
Immunostaining antibodies and reagents and their final concentrations.

**Figure S6.**
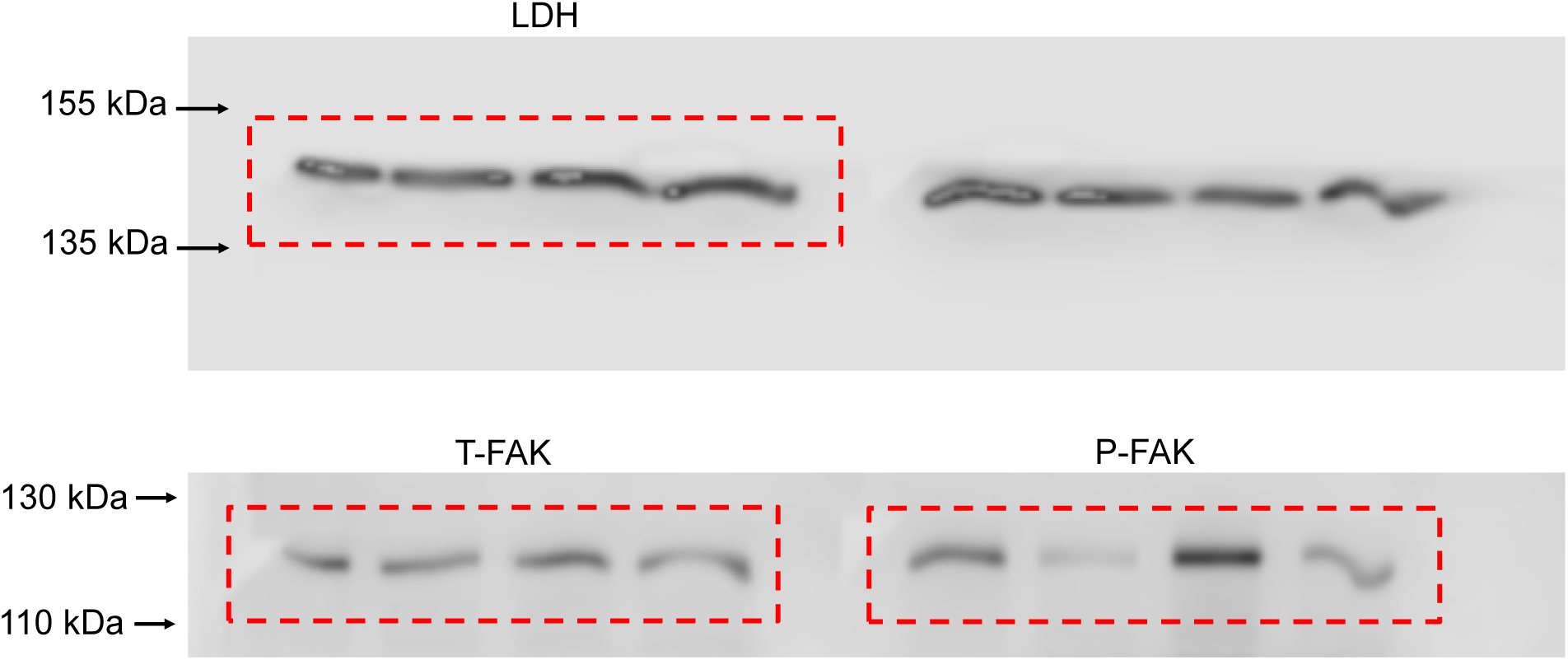
Unprocessed blots used in Figure S3 as obtained by LiCor C-DiGit blot scanner.

## References

1 Smith, S. M. et al. Calcium metabolism before, during, and after a 3-mo spaceflight: kinetic and biochemical changes. Am J Physiol 277, R1–10 (1999).

2 Greenleaf, J. E., Bulbulian, R., Bernauer, E. M., Haskell, W. L. & Moore, T. Exercise-training protocols for astronauts in microgravity. Journal of applied physiology (Bethesda, Md. : 1985) 67, 2191–2204, doi:10.1152/jappl.1989.67.6.2191 (1989).

3 Vico, L. et al. Effects of long-term microgravity exposure on cancellous and cortical weight-bearing bones of cosmonauts. Lancet (London, England) 355, 1607–1611 (2000).

4 Thompson, W. R., Rubin, C. T. & Rubin, J. Mechanical regulation of signaling pathways in bone. Gene 503, 179–193, doi:10.1016/j.gene.2012.04.076 (2012).

5 Ozcivici, E. et al. Mechanical signals as anabolic agents in bone. Nature reviews. Rheumatology 6, 50–59, doi:10.1038/nrrheum.2009.239 (2010).

6 Rando, T. A. & Ambrosio, F. Regenerative Rehabilitation: Applied Biophysics Meets Stem Cell Therapeutics. Cell Stem Cell 22, 306–309, doi:10.1016/j.stem.2018.02.003 (2018).

7 Chan, M. E., Uzer, G. & Rubin, C. The Potential Benefits and Inherent Risks of Vibration as a Non-Drug Therapy for the Prevention and Treatment of Osteoporosis. Current osteoporosis reports, 1–9, doi:10.1007/s11914-012-0132-1 (2013).

8 Judex, S., Gross, T. S. & Zernicke, R. F. Strain Gradients Correlate with Sites of Exercise-Induced Bone-Forming Surfaces in the Adult Skeleton. Journal of Bone and Mineral Research 12, 1737–1745, doi:10.1359/jbmr.1997.12.10.1737 (1997).

9 Rubin, C. T. & Lanyon, L. E. Dynamic strain similarity in vertebrates; an alternative to allometric limb bone scaling. Journal of Theoretical Biology 107, 321–327, doi:10.1016/s0022-5193(84)80031-4 (1984).

10 Price, C., Zhou, X. Z., Li, W. & Wang, L. Y. Real-Time Measurement of Solute Transport Within the Lacunar-Canalicular System of Mechanically Loaded Bone: Direct Evidence for Load-Induced Fluid Flow. Journal of Bone and Mineral Research 26, 277–285, doi:10.1002/jbmr.211 (2011).

11 Gurkan, U. A. & Akkus, O. The Mechanical Environment of Bone Marrow: A Review. Annals of biomedical engineering 36, 1978–1991, doi:10.1007/s10439-008-9577-x (2008).

12 Vainionpaa, A. et al. Intensity of exercise is associated with bone density change in premenopausal women. Osteoporosis international : a journal established as result of cooperation between the European Foundation for Osteoporosis and the National Osteoporosis Foundation of the USA 17, 455–463, doi:10.1007/s00198-005-0005-x (2006).

13 Dickerson, D. A., Sander, E. A. & Nauman, E. A. Modeling the mechanical consequences of vibratory loading in the vertebral body: microscale effects. Biomechanics and Modeling in Mechanobiology 7, 191–202, doi:10.1007/s10237-007-0085-y (2008).

14 Coughlin, T. R. & Niebur, G. L. Fluid shear stress in trabecular bone marrow due to low-magnitude high-frequency vibration. Journal of biomechanics 45, 2222–2229, doi:10.1016/j.jbiomech.2012.06.020 (2012).

15 Riddle, R. C. & Donahue, H. J. From Streaming Potentials to Shear Stress: 25 Years of Bone Cell Mechanotransduction. Journal of Orthopaedic Research 27, 143–149, doi:10.1002/jor.20723 (2009).

16 Fritton, S. P., McLeod, K. J. & Rubin, C. T. Quantifying the strain history of bone: spatial uniformity and self-similarity of low-magnitude strains. Journal of biomechanics 33, 317–325, doi:10.1016/s0021-9290(99)00210-9 (2000).

17 Pagnotti, G. M. et al. Combating osteoporosis and obesity with exercise: leveraging cell mechanosensitivity. Nature Reviews Endocrinology, doi:10.1038/s41574-019-0170-1 (2019).

18 Pongkitwitoon, S., Uzer, G., Rubin, J. & Judex, S. Cytoskeletal Configuration Modulates Mechanically Induced Changes in Mesenchymal Stem Cell Osteogenesis, Morphology, and Stiffness. Scientific reports 6, 34791, doi:10.1038/srep34791 (2016).

19 Uzer, G. et al. Cell Mechanosensitivity to Extremely Low-Magnitude Signals Is Enabled by a LINCed Nucleus. STEM CELLS 33, 2063–2076, doi:10.1002/stem.2004 (2015).

20 Uzer, G., Pongkitwitoon, S., Ete Chan, M. & Judex, S. Vibration induced osteogenic commitment of mesenchymal stem cells is enhanced by cytoskeletal remodeling but not fluid shear. Journal of Biomechanics 46, 2296–2302, doi:10.1016/j.jbiomech.2013.06.008 (2013).

21 Pan, J. X. et al. YAP promotes osteogenesis and suppresses adipogenic differentiation by regulating beta-catenin signaling. Bone research 6, 18, doi:10.1038/s41413-018-0018-7 (2018).

22 Dupont, S. et al. Role of YAP/TAZ in mechanotransduction. Nature 474, 179–183, doi:http://www.nature.com/nature/journal/v474/n7350/abs/10.1038-nature10137-unlocked.html#supplementary-information (2011).

23 Kegelman, C. D. et al. Skeletal cell YAP and TAZ combinatorially promote bone development. FASEB journal : official publication of the Federation of American Societies for Experimental Biology 32, 2706–2721, doi:10.1096/fj.201700872R (2018).

24 Zhao, B. et al. TEAD mediates YAP-dependent gene induction and growth control. Genes & development 22, 1962–1971, doi:10.1101/gad.1664408 (2008).

25 Benham-Pyle, B. W., Pruitt, B. L. & Nelson, W. J. Cell adhesion. Mechanical strain induces E-cadherin-dependent Yap1 and beta-catenin activation to drive cell cycle entry. Science (New York, N.Y.) 348, 1024–1027, doi:10.1126/science.aaa4559 (2015).

26 Ho, L. T. Y., Skiba, N., Ullmer, C. & Rao, P. V. Lysophosphatidic Acid Induces ECM Production via Activation of the Mechanosensitive YAP/TAZ Transcriptional Pathway in Trabecular Meshwork Cells. Investigative Ophthalmology & Visual Science 59, 1969–1984, doi:10.1167/iovs.17-23702 (2018).

27 Cai, H. & Xu, Y. The role of LPA and YAP signaling in long-term migration of human ovarian cancer cells. Cell Communication and Signaling 11, 31, doi:10.1186/1478-811X-11-31 (2013).

28 Driscoll, Tristan P., Cosgrove, Brian D., Heo, S.-J., Shurden, Zach E. & Mauck, Robert L. Cytoskeletal to Nuclear Strain Transfer Regulates YAP Signaling in Mesenchymal Stem Cells. Biophysical journal 108, 2783–2793, doi:10.1016/j.bpj.2015.05.010 (2015).

29 Janmaleki, M., Pachenari, M., Seyedpour, S. M., Shahghadami, R. & Sanati-Nezhad, A. Impact of Simulated Microgravity on Cytoskeleton and Viscoelastic Properties of Endothelial Cell. Scientific reports 6, 32418, doi:10.1038/srep32418 (2016).

30 Qian, A. R. et al. Fractal Dimension as a Measure of Altered Actin Cytoskeleton in MC3T3-E1 Cells Under Simulated Microgravity Using 3-D/2-D Clinostats. Biomedical Engineering, IEEE Transactions on 59, 1374–1380, doi:10.1109/TBME.2012.2187785 (2012).

31 Pardo, S. J. et al. Simulated microgravity using the Random Positioning Machine inhibits differentiation and alters gene expression profiles of 2T3 preosteoblasts. Am J Physiol Cell Physiol 288, doi:10.1152/ajpcell.00222.2004 (2005).

32 Dai, Z. Q., Wang, R., Ling, S. K., Wan, Y. M. & Li, Y. H. Simulated microgravity inhibits the proliferation and osteogenesis of rat bone marrow mesenchymal stem cells. Cell proliferation 40, 671–684, doi:10.1111/j.1365-2184.2007.00461.x (2007).

33 Shi, F. et al. Simulated Microgravity Promotes Angiogenesis through RhoA-Dependent Rearrangement of the Actin Cytoskeleton. Cellular physiology and biochemistry : international journal of experimental cellular physiology, biochemistry, and pharmacology 41, 227–238, doi:10.1159/000456060 (2017).

34 Corydon, T. J. et al. Reduced Expression of Cytoskeletal and Extracellular Matrix Genes in Human Adult Retinal Pigment Epithelium Cells Exposed to Simulated Microgravity. Cellular physiology and biochemistry : international journal of experimental cellular physiology, biochemistry, and pharmacology 40, 1–17, doi:10.1159/000452520 (2016).

35 Chen, Z., Luo, Q., Lin, C., Kuang, D. & Song, G. Simulated microgravity inhibits osteogenic differentiation of mesenchymal stem cells via depolymerizing F-actin to impede TAZ nuclear translocation. Scientific reports 6, 30322, doi:10.1038/srep30322 (2016).

36 Uddin, S. M. & Qin, Y. X. Enhancement of osteogenic differentiation and proliferation in human mesenchymal stem cells by a modified low intensity ultrasound stimulation under simulated microgravity. PLoS One 8, e73914, doi:10.1371/journal.pone.0073914 (2013).

37 Touchstone, H. et al. Recovery of stem cell proliferation by low intensity vibration under simulated microgravity requires intact LINC complex npj. Microgravity 5, doi:doi.org/10.1038/s41526-019-0072-5 (2019).

38 Shiu, J.-Y., Aires, L., Lin, Z. & Vogel, V. Nanopillar force measurements reveal actin-cap-mediated YAP mechanotransduction. Nature cell biology 20, 262–271, doi:10.1038/s41556-017-0030-y (2018).

39 Saeed, M. & Weihs, D. Finite element analysis reveals an important role for cell morphology in response to mechanical compression. Biomechanics and Modeling in Mechanobiology, doi:10.1007/s10237-019-01276-5 (2019).

40 Hu, J. K.-H. et al. An FAK-YAP-mTOR Signaling Axis Regulates Stem Cell-Based Tissue Renewal in Mice. Cell Stem Cell 21, 91-106.e106, doi:10.1016/j.stem.2017.03.023 (2017).

41 Riddick, N., Ohtani, K. & Surks, H. K. Targeting by myosin phosphatase-RhoA interacting protein mediates RhoA/ROCK regulation of myosin phosphatase. Journal of Cellular Biochemistry 103, 1158–1170, doi:10.1002/jcb.21488 (2008).

42 Sen, B. et al. Mechanical signal influence on mesenchymal stem cell fate is enhanced by incorporation of refractory periods into the loading regimen. Journal of biomechanics 44, 593–599, doi:10.1016/j.jbiomech.2010.11.022 (2011).

43 De Magistris, P. & Antonin, W. The Dynamic Nature of the Nuclear Envelope. Current biology : CB 28, R487–r497, doi:10.1016/j.cub.2018.01.073 (2018).

44 Czapiewski, R., Robson, M. I. & Schirmer, E. C. Anchoring a Leviathan: How the Nuclear Membrane Tethers the Genome. Frontiers in genetics 7, 82, doi:10.3389/fgene.2016.00082 (2016).

45 Uzer, G. et al. Sun-mediated mechanical LINC between nucleus and cytoskeleton regulates betacatenin nuclear access. J Biomech 74, 32–40, doi:10.1016/j.jbiomech.2018.04.013 (2018).

